# Using Deep Learning for Gene Detection and Classification in Raw Nanopore Signals

**DOI:** 10.1101/2021.12.23.473143

**Authors:** Marketa Nykrynova, Vojtech Barton, Roman Jakubicek, Matej Bezdicek, Martina Lengerova, Helena Skutkova

**Affiliations:** Department of Biomedical Engineering, Faculty of Electrical Engineering, Brno University of Technology, Brno 61600, Czechia; Department of Internal Medicine – Hematology and Oncology, University Hospital Brno, Brno 62500, Czechia

**Keywords:** Nanopore sequencing, Squiggles, Neural network, MLST, Bacterial typing

## Abstract

Recently, nanopore sequencing has come to the fore as library preparation is rapid and simple, sequencing can be done almost anywhere, and longer reads are obtained than with next-generation sequencing. The main bottleneck still lies in data postprocessing which consists of basecalling, genome assembly, and localizing significant sequences, which is time consuming and computationally demanding, thus prolonging delivery of crucial results for clinical practice. Here, we present a neural network-based method capable of detecting and classifying specific genomic regions already in raw nanopore signals – squiggles. Therefore, the basecalling process can be omitted entirely as the raw signals of significant genes, or intergenic regions can be directly analysed, or if the nucleotide sequences are required, the identified squiggles can be basecalled, preferably to others. The proposed neural network could be included directly in the sequencing run, allowing real-time squiggle processing.

## Introduction

DNA sequencing technologies revolutionized our ability to study genetic variations at the molecular level, which is necessary for a broad spectrum of applications from bacterial typing to cancer research. The introduction of the latest technology - nanopore sequencing - was a major breakthrough [1]. Nanopore sequencing, unlike other methods, does not require DNA synthesis or amplification. Sample preparation is significantly more straightforward, and sequencing can be performed practically anywhere [2]–[4]. The technology uses small pores located in a membrane to which a voltage is applied. When the DNA strand passes through the pore, the electric current’s changes are measured, and as a result, the signal of the read called a ‘squiggle’ is obtained.

Besides the advantages of easy preparation and simple use, nanopore sequencing also provides reads with lengths from tens to hundreds of kilobase pairs (kbp), with a record of more than 2 Mbp [5]. However, the main bottleneck in this technology lies in the low read accuracy, which is improving with each chemistry update, yet still does not compare with second-generation sequencers. Inaccuracies also emerged during the basecalling process, where the squiggles are converted to nucleotide sequences. The basecallers achieve an average read accuracy of 85-95 % [6], which is insufficient for analyses such as single nucleotide variant detection. Although using high genome coverage in post-sequencing analysis, the consensus accuracy can be improved to 99.9 % [7]. If the basecalling process was bypassed and an analysis of the raw squiggles was performed, the results would be more precise and delivered in a shorter time as the Oxford Nanopore Technologies MinION sequencing platform allows real-time access to the sequencing run [8]. Thus, waiting for the whole run to finish would not be necessary, and the crucial information could be analysed almost in real-time. As the squiggle analysis can be performed during the sequencing itself, and the sequencing process can be stopped based on the real-time outputs from the analytical software, there is no need to use a flowcell’s whole sequencing capacity.

As mentioned above, the basecalling process has lower read accuracy caused by several problems. Firstly, the change in current deviation does not correspond to one nucleotide passing through the nanopore, but approximately five nucleotides, leading to 4^5^ = 1024 current levels [9]. Moreover, the number of possible current levels can be higher because the 5-nucleotide step and the speed of the nucleotide strand passing through the nanopore are not constant. Secondly, the same bases can be chemically or epigenetically modified (e.g. 5-methylcytosine) [10], and the whole signal is also affected by noise. These complications mean that only neural networks (NNs) are used for basecalling nowadays [10]. NNs have also found applications in other parts of nanopore sequencing, such as selective sequencing. An example of a tool used to determine whether to eject the sequenced molecule or continue sequencing could be SquiggleNet [11], which uses a convolutional neural network learned from the reference organism’s sequencing data. The classifier then decides the sequenced segment’s location and whether to continue sequencing or eject the molecule. Recently, NNs were also employed to distinguish mitochondrial DNA from genomic DNA in squiggles during a sequencing run [12].

With NNs, it would be possible to identify specific genomic regions in raw nanopore data without the need for basecalling, as shown in the presented article. Even more genes or gene fragments can be found once the neural network can recognize and classify them. The proposed NN was used to predict and classify seven multilocus sequence typing (MLST) loci (*gapA, infB, mdh, pgi, phoE, rpoB, tonB*) in 29 *Klebsiella pneumoniae* genome squiggles. Raw squiggle analysis can bring more precise information, as the epidemiological and chemical modification can also be studied. In the future, distinguishing bacterial strains could be done using only the signals with no basecalling, providing crucial epidemiological information for early outbreak identification.

## Results

### Analysed genes

The seven MLST loci (*gapA, infB, mdh, pgi, phoE, rpoB, tonB*) from BIGSdb [13] were chosen for showing the proposed neural network’s performance. The first allele from each MLST gene was searched for in basecalled reads. The median lengths of the located MLST genes were, on average, 4 bp shorter than the queries. Thus, most of the identified genes contained almost the whole allele sequence. The analysed sequence variability was about 6.31% (**Table 1)**. This variability is caused mainly by the presence of other alleles from a given gene in the dataset, as only one allele for each gene was searched for. Based on these values, it can be said that the dataset is variable enough to train the network to be able to recognize different gene’s alleles.

**Table 1.**
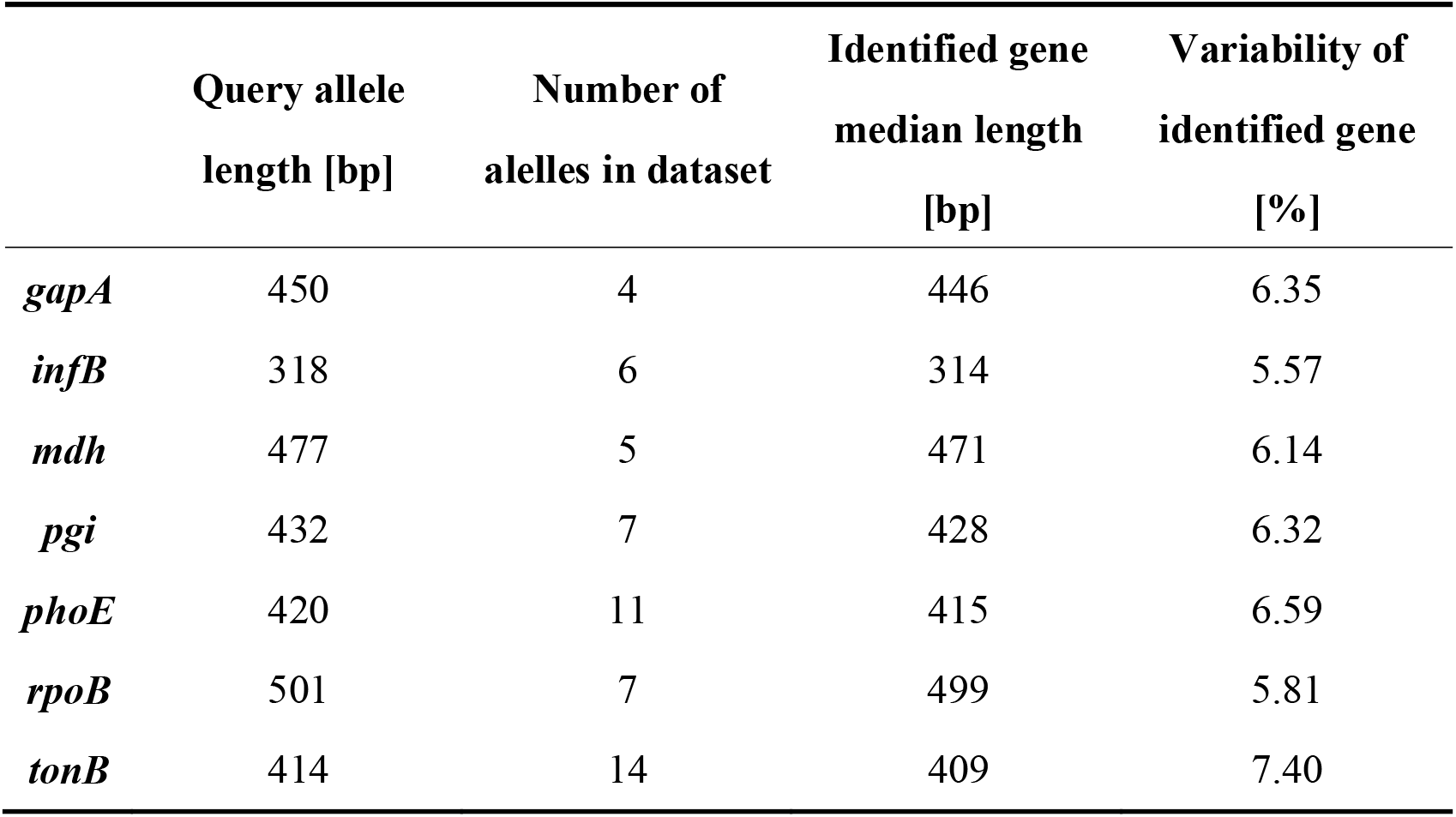
Statistics about seven analysed MLST loci.

### Dataset preparation

Gene signals that could be used to train the neural network and later validate its performance were obtained by the following described process. BLAST (2.9.0+, [14]) was employed to localize sequences of interest. The fast5 files containing basecalled fastq sequences were used to prepare the signal database. Then, the gene templates were examined in basecalled data, and results (hits) were saved in CSV format. In addition, the whole squiggles containing the gene sequences were extracted and saved in an internal h5 format, so they could be swiftly accessible and easily modified.

If more genomes were sequenced in one run, the BLAST results contained hits for all genomes from a particular run. To assign barcodes to the hits, demultiplexing tables were created for sequencing runs where barcoding kits were used. The outputs from the Guppy barcoder were used for this purpose. Each created table contained all reads ID from the sequencing run and their corresponding barcodes; so, it was possible to add sample identification to the hits in the BLAST table.

The BLAST results were filtered to remove random and partial hits. For further processing, only the hits with a percentage of identical matches more significant or equal to 90 %, the length at least 90 % of query length, and the e-value lower or equal to 1e-50 were chosen. The searched sequences were found on both the leading and complementary strands.

In the last signal extraction step, the gene sequences’ BLAST coordinates were recalculated to signal coordinates. Thus, a dataset containing squiggles with desired genes was created. To each squiggle, the gene signal coordinates were added. The neural network should also recognize squiggles without genes; therefore, datasets with no genes were created.

For preprocessing the raw sequencing fast5 files and dataset preparation, the internally developed MANASIG [15] package was used and is available on <u>GitHub</u>.

### Signals for training/validating neural network

In total, 48,860 squiggles from which 38,867 contained one of the seven housekeeping genes fragments were analysed. The length of squiggles with genes ranged from 4,362 to 4,477,607 samples with a median of about 142,869 samples. The median gene fragment lengths were 4,815 (*gapA*), 3,417 (*infB*), 5,197 (*mdh*), 4,643 (*pgi*), 4,586 (*phoE*), 5,387 (*rpoB*) and 4,565 (*tonB*) samples. The number of signals with no genes was 9,993 and their lengths varied from 2,008 to 1,411,573 samples with a median value of 24,408 samples. These squiggles were randomly generated from the seven sequencing runs. For a detailed number of squiggles from each analysed genome, see Table S1.

For neural network training, about 75 % of all squiggles were used, and the rest were used to validate NN performance.

### Gene detection in squiggles

From the validation dataset, 41 squiggles with corrupted gene coordinates were removed. In total, 11,887 squiggles were used for validation, of which 2,000 had no gene sequences. The representation of individual genes in the dataset was as follows: *gapA* – 1,671, *infB* – 1,570, *mdh* – 1,644, *pgi* – 1,659, *phoE* – 1,160, *rpoB* – 1,179 and *tonB* – 1,004.

The gene predictions in the downsampled squiggles containing any genes of interest were classified into two categories – a gene found, and a gene not found. To evaluate, the calculated dice coefficients were used. On average, the detection success rate was about 98%; see **Table 2** for specific values. The gene found category was further split into the other two subgroups – the gene was found correctly (dice ≥ 0.9), or the gene was shifted. If the predicted gene was labelled as shifted, four variants could be observed – the detected part was inside/outside the annotated borders or shifted before/beyond the start/end of the gene. Examples of all possible gene detection cases are shown in **Figure 1**, and squiggle percentages in the given categories can be seen in **Table 3**. The prediction with dice greater than or equal to 0.9 differed from the annotation position by, on average, about ten samples. Predictions with lower dice consisted mainly of genes shifted beyond the annotated gene start and end, and from predictions inside annotated gene boundaries.

**Table 2.**
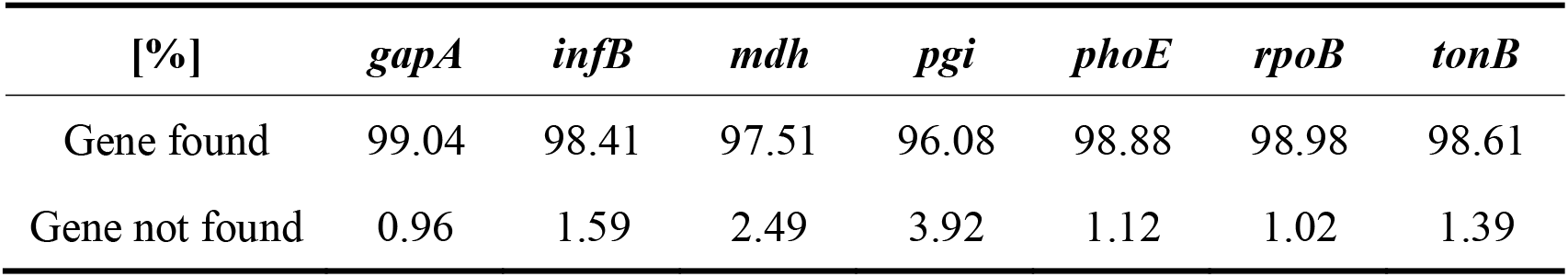
Percentages of identified or non-identified MLST loci.

**Table 3.**
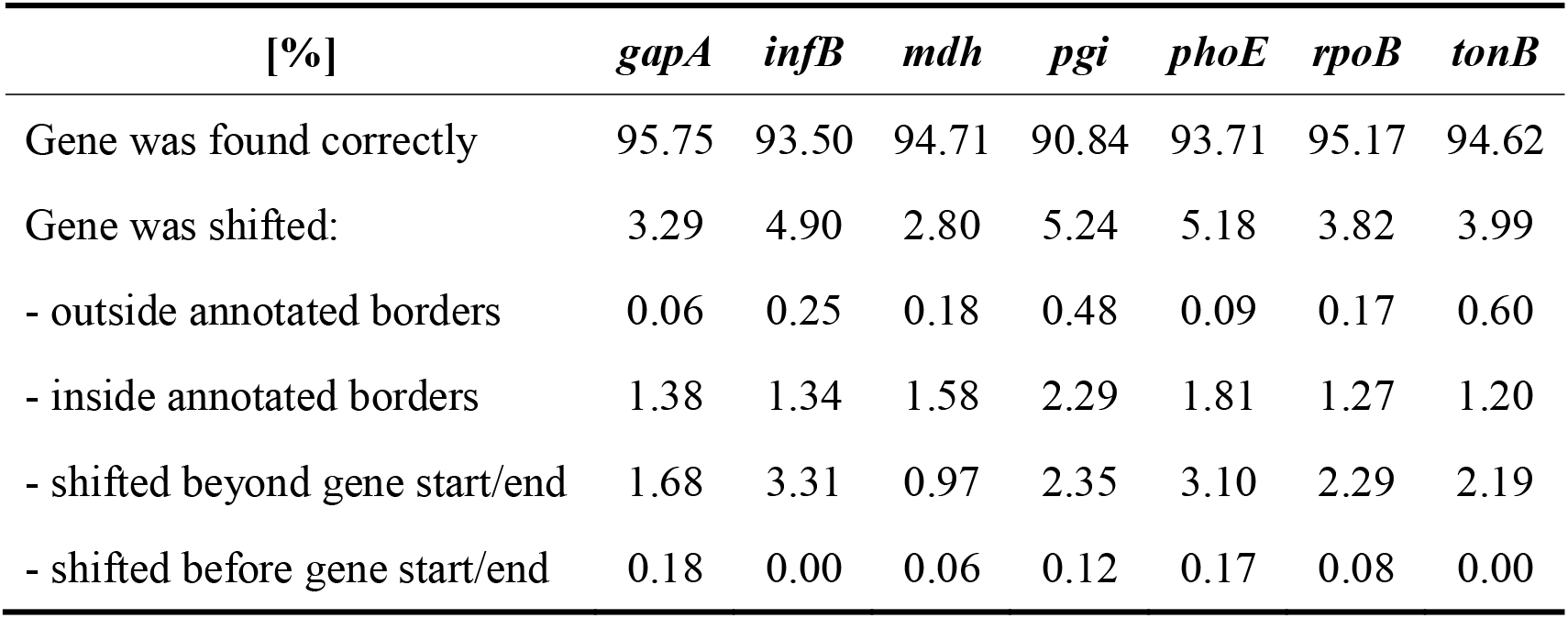
Percentages of identified MLST loci including shifted genes.

| [%] | *gapA* | *infB* | *mdh* | *pgi* | *phoE* | *rpoB* | *tonB* |
| --- | --- | --- | --- | --- | --- | --- | --- |
| Gene was found correctly | 95.75 | 93.50 | 94.71 | 90.84 | 93.71 | 95.17 | 94.62 |
| Gene was shifted: | 3.29 | 4.90 | 2.80 | 5.24 | 5.18 | 3.82 | 3.99 |
| - outside annotated borders | 0.06 | 0.25 | 0.18 | 0.48 | 0.09 | 0.17 | 0.60 |
| - inside annotated borders | 1.38 | 1.34 | 1.58 | 2.29 | 1.81 | 1.27 | 1.20 |
| - shifted beyond gene start/end | 1.68 | 3.31 | 0.97 | 2.35 | 3.10 | 2.29 | 2.19 |
| - shifted before gene start/end | 0.18 | 0.00 | 0.06 | 0.12 | 0.17 | 0.08 | 0.00 |

**Figure 1.**
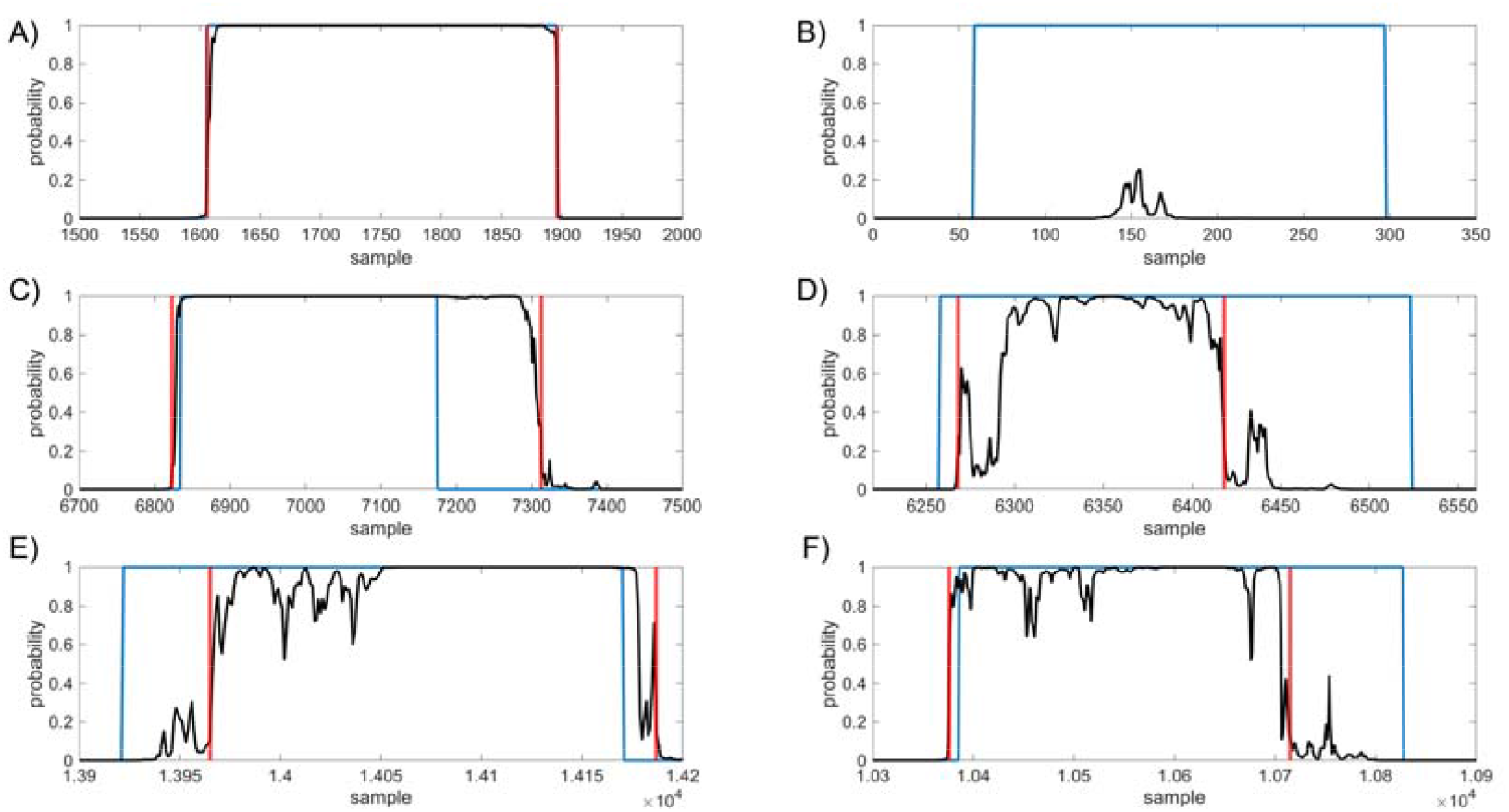
Examples of gene detection, annotated positions are blue, gene occurrence functions are black and detected gene positions are red. A. gene was found correctly, B. gene was not found, C. gene was outside annotated borders, D. gene was inside annotated borders, E. gene was shifted to right, F. gene was shifted to left.

In the case of squiggles with no genes, predictions with dice equal to one were marked as correct. In total, 96.6% of squiggles without any genes were successfully recognized, and no gene was detected in them.

### Gene squiggles classification

The squiggles from the validation dataset were classified via the proposed neural network into eight categories, where seven categories were for analysed genes, and the last one was for the squiggles with no gene. For each squiggle, the probabilities that the gene belongs to a given category were established. From these values, the maximum was chosen, and the squiggles were assigned to a given category.

The squiggles with no MLST genes were correctly classified in 99.80% of cases and only four squiggles were misclassified. The percentage of successful gene classification was about 94.67% for five out of seven MLST loci. In the case of *pgi*, the success rate dropped to 87.64% and in the case of *mdh* to 83.33%. See the detailed results in **Figure 2**.

**Figure 2.**
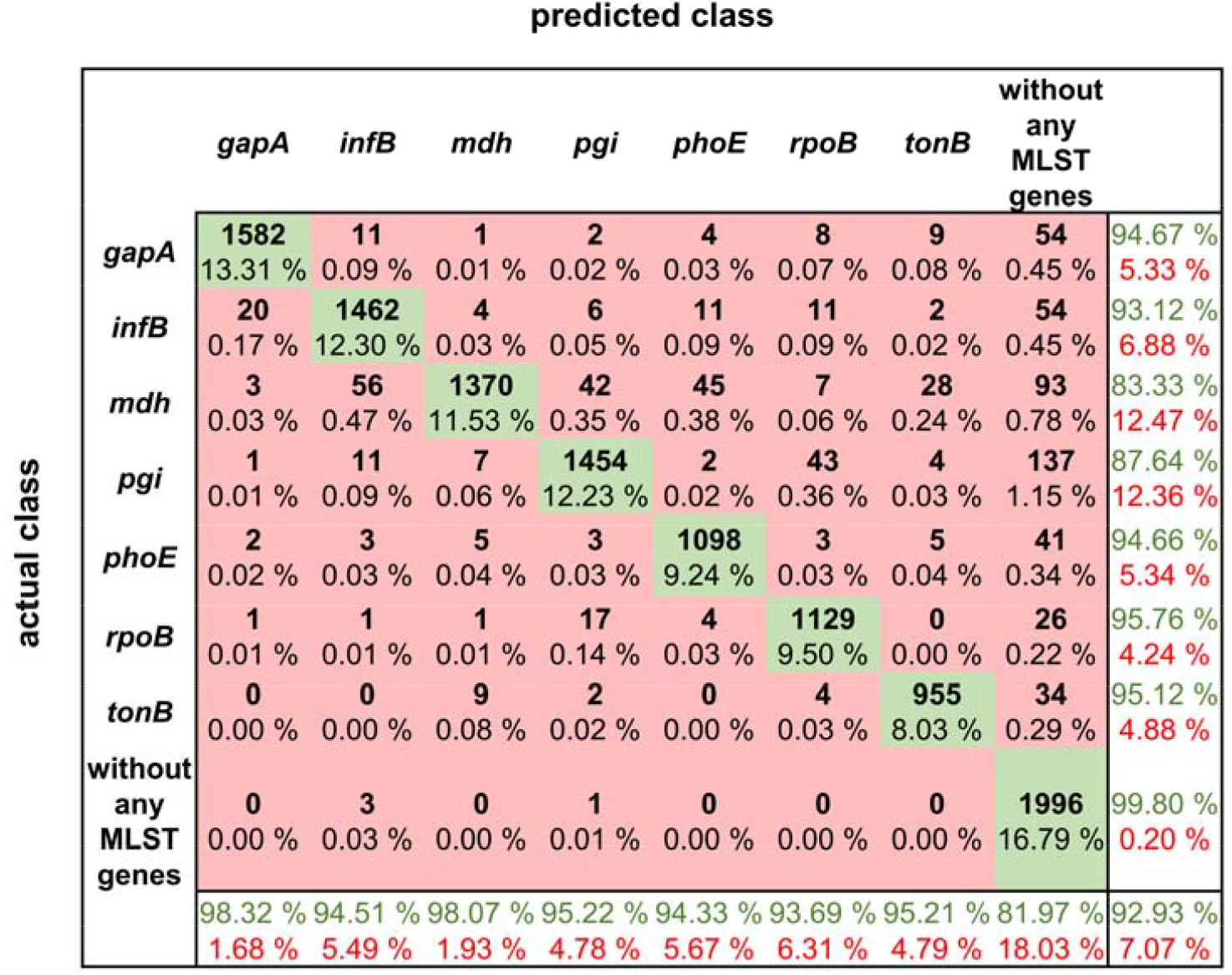
Confusion matrix for MLST loci classification results.

The confusion matrix shows the true positive rate, which is the highest for the squiggles with no detectable gene and in the case of MLST genes, the highest value of 95.76% is for *rpoB* gene. In general, the normalized true positive value for all MLST genes was 92.04%; if squiggles without genes are included, the average true positive rate is even higher, at 93.01%.

On the other hand, the false discovery rate was the highest for the squiggles without genes. The difference between the average false discovery rate for the squiggles with and without genes is 13.65%. It can be concluded that there is a much higher rate of not identifying the gene than identifying it incorrectly.

## Discussion

A comparison of detected and annotated gene coordinates (**Figure 3**) and their overlaps was conducted. The results showed that they came from the same statistical distribution. The neural network tended to identify the genes a bit longer than annotated. This phenomenon can be caused by the signal length irregularity in the same sequence. Also, the signals were downsampled for the neural network. This pre-processing and the need to recompute the positions of genes to the original space signal can cause some variations in the genes’ signals lengths.

**Figure 3.**
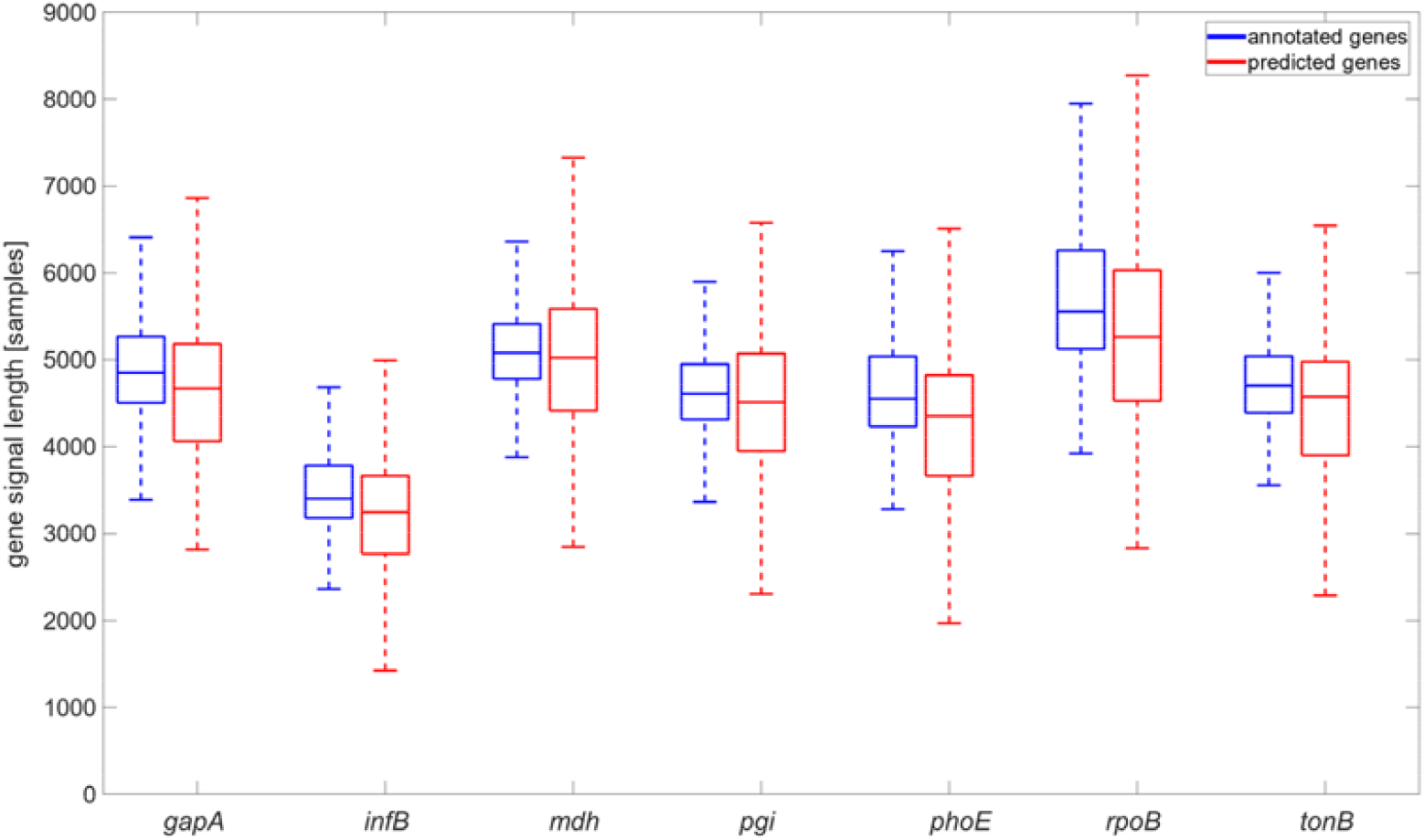
Comparison of annotated gene signal length and gene signal length identified by the neural network, graph is shown without flyers for clearness.

The sequencing data are stored in the fast5 format, which is based on hdf5 format. However, hdf5 is not fully backwards compatible; the hierarchical structure can change with every new nanopore chemistry or software upgrade, and no complex descriptions about the fast5 nanopore format and its modification exist. For this reason, our proposed package for fast5 file processing can be used with R9.4.1 flowcells, two sequencing kits (SQK-RBK004, SQK-LSK109) and nanopore MinKNOW v19 and v20 sequencing software; with other versions, it might be necessary to modify the package. Completely new chemistry can also influence signal properties such as squiggle lengths, and in that case, new neural network training would be needed. Also, the neural network may not work correctly if the squiggle lengths in the training/validating dataset contain outliers such as extremely long ones.

During the gene coordinates’ detection in predicted signals, it was found that gene occurrence functions sometimes contained more than just one peak exceeding a specified threshold, as shown in **Figure 4A** and **Figure 4B**. Multiple predictions were observed in 3.81% squiggles from the validation dataset, and in the majority of cases (more than 90%), two peaks were predicted. After detecting peak coordinates, the corresponding basecalled nucleotide sequences were analysed. It was found out that the false positive predictions contained sequences with a partial match to the desired gene, see **Figure 4C**. Nevertheless, the partial matches were significantly shorter than the examined sequences; therefore, they could be filtered in postprocessing. However, there could be a problem with setting the filter parameters because the partial matches may have a different numbers of samples in the squiggles as the speed of DNA passing through the pore is not constant and the sampling is non-equidistant. This mentioned multiple detection problem could be solved if the genes or other specific sequences we wanted to search for are unique and non-repetitive in the analysed bacterial species. On the other hand, from multiple detection results, it can also be said that the neural network could find even short signals that corresponded to several dozen base pair long sequences, such as parts of genes. In addition, it could recognize the squiggles even if there were many mutations; thus, the network can be used to predict and classify even highly variable genomic regions.

**Figure 4.**
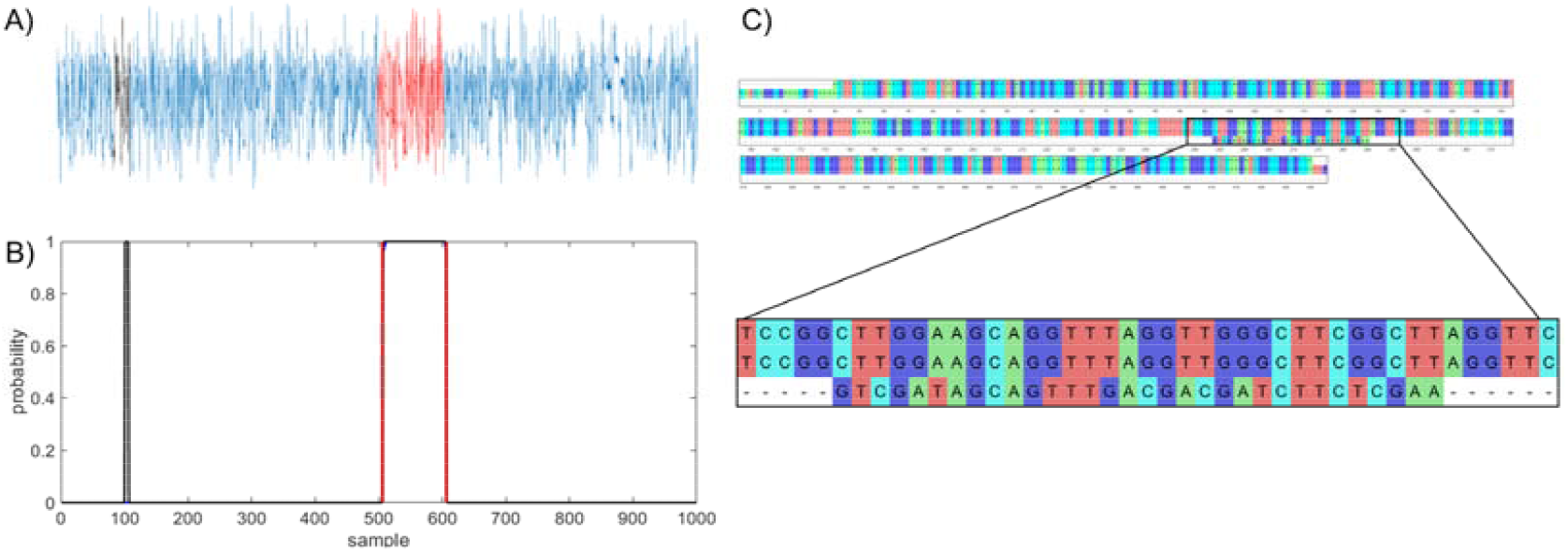
Example of multiple gene prediction. A. squiggle with labelled positions of gene (red) and random match (black), B. graph of predicted gene in squiggle, gene occurrence function is black, annotated position is red, C. multiple sequence alignment of *gapA* template, *gapA* hit and random match.

If more than one gene is located in the analysed squiggle, it could cause a problem for proper neural network function. Multiple gene presence may occur if long squiggles are produced during the sequencing process. To avoid this problem, genomic regions we wanted to detect should be carefully selected. The first option is to choose genes that have sufficient distance from each other in the analysed species. The second option is to pre-process the squiggles before sending them into a neural network. Signals longer than a given threshold can be divided into several shorter signals, ensuring that only one targeted region will be present in each signal.

## Conclusion

This paper presented a deep learning method to identify and classify specific genomic regions in raw nanopore sequencing data. The proposed neural network can be used to find whole genes, their parts or intergenic regions. We showed one of the possible neural network uses – detecting and classifying seven MLST loci in *K. pneumoniae* genomes in squiggles.

The percentage of correctly predicted genes was 98.2%, and they were successfully classified in 92.9% cases. The squiggles with no MLST loci were correctly predicted in 96.6% and classified in 99.8% cases. The NN achieved the same accuracy as basecalling tools, and if postprocessing was employed, the accuracy could be even higher.

The main requirement of the proposed approach for gene prediction and classification is a large amount of data to train the neural network. Without sufficient data, the network would not be adequately trained, and precise results could not be obtained. From the study, it can also be said that the genomic regions detected via the neural network should be unique, non-repetitive sequences of any length and should be located in the analysed genomes with sufficient distances between them.

Nanopore sequencing has huge potential in routine clinical practice. It can be used instead of time consuming NGS to deliver crucial epidemiology information earlier. For example, if there is a need to analyse samples from one hospital department to find out if there is an outbreak or not, if an infection is spreading via instruments, patients or medical staff, nanopore sequencing can be employed. NGS analysis would consist of library preparation, sequencing, and in silico sequence type determination from post-processed sequencing data, which can take two to three days while the potential epidemic spreads uncontrollably. The results could be delivered the same morning if nanopore sequencing combined with the proposed NN is employed. The squiggles containing MLST loci can be recognized during a run or immediately after it and analysed. If more genomes are sequenced in one run, the barcodes attached to MLST squiggles can be used to find out which, e.g. *gapA* loci belongs to which genome.

The network can be trained to recognize different significant sequences. Hence, the proposed approach can be used for other purposes, such as direct positive clinical sample sequencing. The network can filter out the human sequences, identify MLST loci, and determine infectious agents’ sequence types. The cultivation will not be needed, which significantly saves time.

## Materials and methods

### Analyzed bacterium

*K. pneumoniae* is a Gram-negative opportunistic pathogen from the *Enterobacteriaceae* family. Usually, it affects immunocompromised patients, and the majority of *K. pneumoniae* infections are hospital-acquired. The gastrointestinal tract colonization generally occurs before nosocomial infections develop, which usually affect the urinary tract, respiratory tract or result in septicemia or soft tissue infection [16],[17]. The genome size of *K. pneumoniae* is about 5.5 Mbp and incorporates about 5,000 to 6,000 genes from which 2,000 genes form the core genome, and almost 30,000 genes are parts of the pangenome [18].

### Genome sequencing

In this article, 29 *K. pneumoniae* isolates collected between 09/2014 and 07/2019, mainly at the Department of Internal Medicine, Hematology and Oncology at the University Hospital Brno, were analysed. The high molecular weight DNA was extracted using the MagAttract HMW DNAKit (Qiagen, Venlo, NL), and the NanoDrop (Thermo Fisher Scientific, Waltham, MA, USA) was employed to measure the purity of the extracted DNA. The DNA concentration was checked by Qubit 3.0 Fluorometer (Thermo Fisher Scientific, Wilmington, DE, United States) and using Agilent 4200 TapeStation (Agilent Technologies, Santa Clara, CA, USA) the proper length of the isolated DNA was checked. The Rapid Barcoding Kit (Oxford Nanopore Technologies, Oxford, UK) was used to prepare the sequencing library for 27 *K. pneumoniae* isolates. For the remaining two samples, the Ligation Sequencing 1D Kit (Oxford Nanopore Technologies, Oxford, UK) was used to prepare the library for Oxford Nanopore sequencing. The sequencing was performed using the MinION sequencing platform (Oxford Nanopore Technologies, Oxford, UK) with R9.4.1 flowcells.

The sequenced genomes were basecalled and, in the case of the pooled library, separated according to barcodes using Guppy software (3.4.4+a296acb). The data quality was checked using PycoQC (v2.2.3, [19]). See Table S2 for detailed information about each sequencing run. The analyzed datasets can be found in the National Center for Biotechnology Information Sequence Read Archive database under a BioProject with accession number <u>PRJNA786743</u>.

### Neural network architecture

To sort out the task of enabling gene localization and classification, some challenges had to be overcome. The main problem was squiggles that were too long, moreover with uneven sampling time. This distinctly increases computational complexity and memory demands, and first and foremost, causes a significant vanishing gradient (VG) effect during training, especially for the recurrent nets. Further, conventional neural networks cannot be applied to signals of different lengths, and they require strictly the same input length.

The proposed NanoGeneNet can be divided into three basic parts: feature extraction, gene localization (so-called a sequence-to-sequence regime) and finally, gene classification with another feature extraction (a sequence-to-vector regime). Using the combination of convolution and recurrent networks turned out to be a great solution for long and uneven signal lengths. For more details on architecture design, see **Figure 5**.

**Figure 5.**
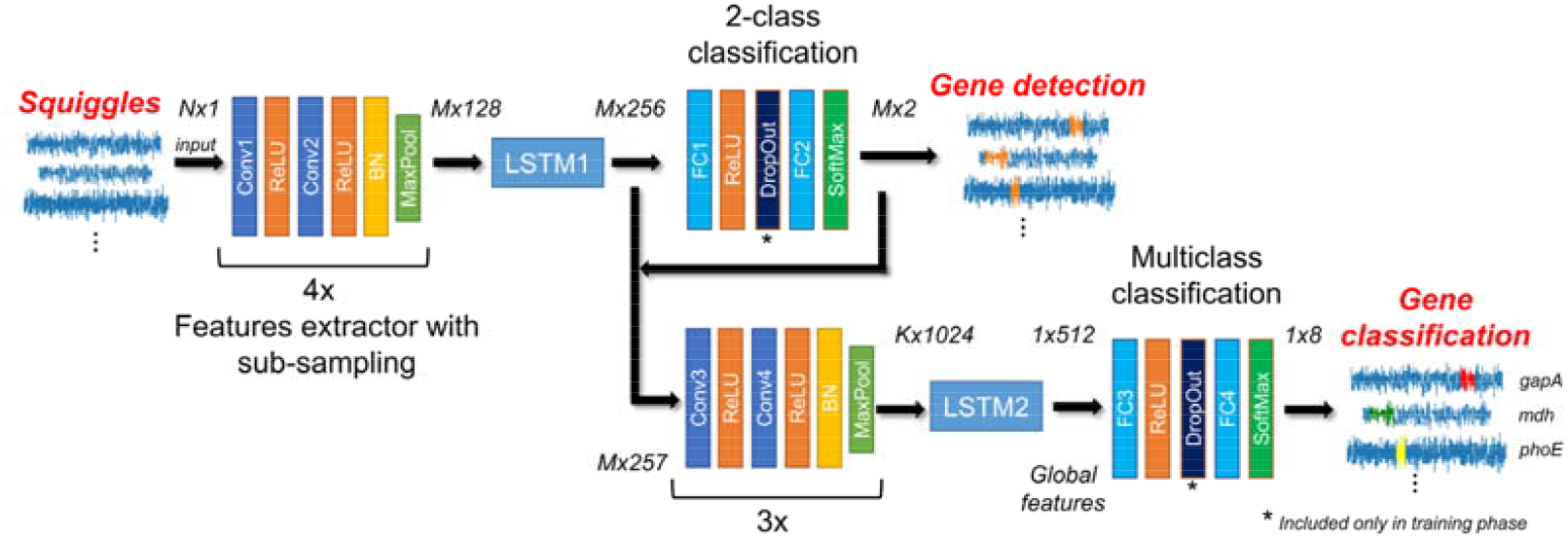
Architecture of NanoGeneNet. Each encoder block contains two convolution layers (Conv) with a kernel size equal to 3, followed by ReLU activation function and finally a Batch Normalization (BN) and a Maxpool layer with size kernel and stride of 2. Network also includes two Long Short-Term Memory (LSTM) recurrent networks and classification blocks composed of Fully Connected layers (FC), SoftMax activation layers and a DropOut layer used only in the training phase. For each block, tensor size shown in the form of *squiggle length x feature number*, where *N* is original squiggle length and *M* is its downsampled version length.

The network was realized in Python 3.9 with the PyTorch library. The source code for training, validation, and especially feedforward for NanoGeneNet is available on <u>GitHub</u> along with a demo and an example squiggle.

#### Local feature extraction with downsampling

The encoder part of the net allows local features to be extracted from the raw signal using a recursive repeating block. As shown in **Figure 5**, the block contains two convolution layers (Conv) with a kernel size of 3 and depth of 16, doubled in each recursion. The Conv layer is always followed by a ReLU activation function, and the block ends with a batch normalization layer and nonlinear spatial reduction layer - MaxPooling. This encoder part occurs twice in the NanoGeneNet.

The network part designed in this way provides a new local feature signal reflecting the occurring or non-occurring gene from an input signal. Due to the MaxPooling layer, there is a spatial reduction in the signal length of *M* with a sub-sampling factor of 2^4^ to decrease computational complexity, while the only relevant features for gene detection are retained during net feedforward. Since this part of the network only views a very local part of the signal, the following LSTM (Long Short-Term Memory) [20] network has the task of viewing the whole signal globally and determining the local features for each signal sample, considering all previous samples (long memory). The output is then the tensor of local feature signals sized *Mx256*, where *M* is the length of the downsampled feature signals.

Due to a recurrent LSTM net, the input can be different length signals and sub-sampling mildly suppresses the VG effect and significantly decreases GPU memory demands. The complete suppression of VG problem is performed by multistage training.

#### Gene occurrence prediction

From the previous LSTM block, each downsampled output signal sample is now coded by feature vector, which inputs into a two-class classification part. Here, two fully connected (FC) networks with ReLU and Softmax activation function can be used, respectively. This part of the proposed NanoGeneNet classifies each downsampled signal sample into two classes; gene or non-gene. The output is tensor *Mx2* defining probability assignment to all classes. Since there are two classes, we can use the probabilities of only the first class as an output gene occurrence function. Examples of such functions are shown in **Figure 1**.

#### Gene classification

In this part, the local feature tensor from the previous encoder and LSTM is concatenated with the obtained likelihood function and the resulting tensor is *Mx257* in size. The tensor is encoded into a new feature tensor corresponding with the feature import to classify gene types via a second encoder block with a sub-sampling factor of 2^3. Then using another LSTM network, the new local feature tensor is transformed into another new tensor (*1×512*) reflecting only the whole signal’s global features to enable classification of the whole signal into gene class. This task is performed via the last part of NanoGeneNet - the multiclass classification network. It has the same architecture as the previous 2-class net, except that the last FC layer has eight neurons followed by SoftMax, enabling a one-hot coding of output class probabilities. The final decision is based on the choice of class with the maximal value.

### Network training details

#### Training and validation dataset

The available signal dataset was divided on the genome level by hand to ensure data distribution consistency for training and validation. For the validation dataset of each gene, all randomly selected genome signals were selected to make up something between 20 % and 30 % of the entire database. It cannot be done exactly because each file contains a different number of signals for specific genes within each genome. As the genome includes the same gene sequence, all signals always within the genome were used for training/testing; ignoring the uneven time sampling, i.e., a nonlinear scaling, which is taken as a kind of augmentation useful for training.

#### Multistage approach

Generally, the VG problem is related to the signal length, the longer the signal, the greater the effect of this problem. Therefore, multistage training was proposed. In the first phase, the first encoder with sub-sampling was pre-trained on shorter signals. In each iteration, the current signal was randomly cut with random termination and random length in a range of 20-80 thousand samples. In the case of gene signal occurrence, this cut training signal always contained the whole gene. In this way, the encoder and LSTM1 training were achieved with a minimal VG problem. To derive a loss function during training, the 2-class classification was trained concurrently based on cut annotations in this way.

In the second phase, the first encoder was additionally retrained on the database containing the non-gene signals. Finally, in this way, the pre-trained net (encoder, LSTM1 and 2-class net) was fine-tuned for signals with genuine length. The same training strategy was also used for the second part of NanoGeneNet, but in this case for an eight-class whole-signal classification task and with its first part frozen.

#### Training parameters

All training dataset signals were randomly shuffled within each new iteration, and all nets were trained from scratch. As both tasks were defined as a classification with a class probability estimation, a weighted cross-entropy was chosen as a loss function. Further, an Adam algorithm (Adaptive Moment Estimation) [21] was used as an optimization algorithm, and the initialization Learning Rate (LR) was set to 0.001, and the weight decay to 0.0001. During all stages of training, the LR was manually changed (decreased) based on the designer’s experiences. A batch size greater than one could only be used in the first training stage, where the net worked with cut signals; here, the batch size was 8, and a larger batch size resulted in too much regularization. In the training phase, the Dropout layer [22] worked with a probability of 50%. Other hyperparameters were set by default in PyTorch, which can be found in the shared source code on <u>GitHub</u>.

#### Hardware requirements

NanoGeneNet training was performed on a computational device with Intel Xeon E5-2603v4, 16 GB RAM and nVidia Titan Xp, 12 GB GDDR5 graphics card. The network was realized in Python 3.9 with the PyTorch library. One stage of training took around two hours, and required up to 10 epochs for a sufficient success rate.

The minimal hardware requirements depended on the input squiggle length, where the above-mentioned device had sufficient parameters to process our database. The prediction and classification of a squiggle with median of length (143 thousand samples) took around 0.2 sec in total. There is a linear dependence between the signal length and computational time, where the longest squiggle from our database (almost 4.5 million samples) took around four seconds.

### Gene coordinates detection in predicted signals

The outputs from the first part were gene occurrence functions determining the probability for each sample in the downsampling signal being part of the gene. The positions of the predicted gene were detected to evaluate the prediction accuracy. In each gene occurrence curve, the peaks exceeding a threshold of 0.9 were found and further analysed as possible target genes. The shortest possible distance between two peaks was set to 100 samples; otherwise, the peaks were merged into one. Boundaries where the gene occurrence curve exceeded 0.5 were located around possible gene peaks in the next step. Then the gene boundaries were extended sample by sample to find a precise prediction beginning and end. The samples were added to the gene until the difference between the two adjacent samples was higher than 0.1.

For each calculated gene’s coordinates, the dice coefficient was calculated as DSC=2TP/(2TP+FP+FN), where *TP* was the number of samples correctly classified as gene, the *FP* was the number of samples falsely labelled as gene, and *FN* was the number of samples falsely labelled as not-gene. If just one peak was observed in the signal, the dice was calculated for it. In cases where more peaks were detected in the gene occurrence function, the coefficient was calculated for each of them. For gene prediction and detection evaluation, the detected gene positions with the highest dice were selected.

## Authors’ contributions

MN, VB, and HS contributed to the conception and design of the study. MN and VB created and implemented the algorithm for fast5 processing and evaluated the results. RJ designed and implemented the neural network. ML and MB ensured the biological aspects of the project. MN and RJ wrote the manuscript. All authors read and approved the final manuscript.

## Acknowledgments

This work has been supported by grant FEKT-K-21-6912 realised within the project Quality Internal Grants of BUT (KInG BUT), Reg. No. CZ.02.2.69 / 0.0 / 0.0 / 19_073 /0016948, which is financed from the OP RDE. Collecting, processing, storing and sequencing of all bacterial isolates used in this study was supported by the Ministry of Health of the Czech Republic, Grant No. NV19-09-00430, all rights reserved.

## Supplementary material

**Table S1 Number of squiggles containing given MLST locus for each analysed genome and number of squiggles with no MLST genes for given sequencing run**

**Table S2 Sequencing runs information**

